# Subgenome-Level Phylogeny Reveals an Allopolyploid Origin of Chloranthales from Ancestral Monocot and Eudicot Lineages

**DOI:** 10.64898/2026.08.22.746209

**Authors:** Yu Cao, Heng-Chi Chen, Yves Van de Peer, Zhen Li, Da-Yong Zhang

## Abstract

Angiosperms (flowering plants) represent the most species-rich lineage of land plants, yet deep phylogenetic relationships among their major clades, particularly within mesangiosperms, remain notoriously difficult to resolve. Although hybridization and whole-genome duplication (WGD) are recognized as major evolutionary forces, the contribution of ancient allopolyploidization, which combines hybridization with WGD, to deep-node phylogenetic discordance in mesangiosperms remains poorly understood.

Here, we analyzed genomes from seven representative early-diverging angiosperm lineages, each containing no more than one lineage-specific WGD event. Using a comparative phylogenomic framework, we characterized WGD-derived paralogs across five WGD-bearing lineages. In contrast to other polyploid lineages, Chloranthales WGD-derived paralogs rarely formed sister relationships in gene trees. After evaluating alternative explanations, including incomplete lineage sorting, ancient paralogy, and analytical artifacts, we demonstrate that this discordant phylogenetic signal reflects the genomic legacy of an ancient allopolyploidization event. We further resolved the Chloranthales subgenomes by exploiting biased fractionation patterns within conserved syntenic regions. Subgenome-resolved phylogenomic analyses revealed that the dominant Chloranthales subgenome exhibits phylogenetic affinity with eudicots, whereas the recessive subgenome is associated with monocots, indicating a deep reticulate origin involving ancestral lineages related to these major angiosperm clades.

Our findings provide subgenome-resolved evidence that ancient allopolyploidization can connect deeply diverged lineages during early angiosperm evolution, highlighting the importance of incorporating reticulate polyploid histories into phylogenomic frameworks for resolving deep evolutionary relationships.

## Introduction

Comprising approximately 90% of all land plant species, flowering plants (angiosperms) dominate terrestrial ecosystems and exhibit remarkable morphological and ecological diversity (1, 2). Over the past two decades, advances in phylogenomics have established that Amborellales, Nymphaeales, and Austrobaileyales (the ANA grade) form successive sister lineages to the remaining angiosperms, collectively termed mesangiosperms (3). Despite this broad consensus, resolving the deep evolutionary relationships among major angiosperm lineages remains highly challenging. Specifically, debates persist over the root of angiosperms (4–6) and the relationships among the five major mesangiosperm lineages (Chloranthales, magnoliids, monocots, Ceratophyllales, and eudicots), for which at least seven distinct topologies have been proposed (7). These conflicting hypotheses have often been attributed to methodological differences, including the choice of genomic markers (nuclear versus organellar genes) and phylogenetic inference strategies (concatenation versus coalescent-based approaches) (3, 6, 8–18). Beyond analytical uncertainties such as phylogenetic estimation error (19), discordance at deep evolutionary nodes is increasingly recognized as a consequence of complex biological processes. These include incomplete lineage sorting (ILS) during the rapid early angiosperm radiations (12, 17), differential gene loss following gene and whole-genome duplications (WGDs) (20), and ancient hybridization or introgression among early diverged lineages (9, 11).

Hybridization and WGD are increasingly recognized as primary drivers of plant speciation, adaptation, and key evolutionary innovations (21–26). Phylogenomic studies have demonstrated that ancient hybridization and introgression are widespread throughout plant evolutionary history and have frequently contributed to the formation of stable and diverse lineages (23). Furthermore, all extant angiosperms are descended from an ancient polyploid ancestor (27–31), and with the potential exceptions of *Amborella* and *Aristolochia*, nearly all sequenced early-diverging lineages have experienced at least one additional, lineage-specific WGD event since diverging from the common ancestor of angiosperms (32, 33). Given the pervasive occurrence of both processes throughout plant evolution, hybridization and WGD likely acted in concert to shape the diversification and genomic architecture of angiosperms. Among polyploid events, allopolyploidization, which combines interspecific hybridization with WGD, introduces complex reticulate signals. Such reticulate origins have been documented across diverse plant lineages, including bamboo (34), poppy (35), cotton (36), poplars and willows (37), and ferns (Thelypteridaceae) (38). Beyond plants, ancient allopolyploidy has been shown to substantially confound phylogenetic relationships among deeply diverged lineages, as exemplified by *Saccharomyces* yeast (39). Nevertheless, despite its potential importance, the contribution of ancient allopolyploidization to phylogenetic discordance among early-diverging angiosperm lineages remains largely unexplored.

Resolving the phylogenetic consequences of allopolyploidization requires subgenome-resolved phylogenies, which depend critically on accurate assignment of genomic regions to their ancestral subgenomes. However, phasing ancient, highly rediploidized paleopolyploid genomes is exceptionally challenging due to extensive chromosomal rearrangements and the lack of extant diploid progenitors (40–42). Phylogeny-based phasing methods are often rendered invalid when progenitors are extinct or unsampled, and can be easily confounded by ILS (43, 44). Alternatively, synteny-based approaches exploit signatures of subgenome dominance caused by biased fractionation, whereby one subgenome preferentially retains more genes and may experience stronger selective constraint than its homoeologous counterpart (45–47). Although fractionation-based approaches do not require extant progenitors, their resolution can be compromised by homoeologous exchange and gene conversion events (42). Therefore, integrating phylogenetic evidence with fractionation-based subgenome inference provides a complementary framework for reconstructing ancient allopolyploid evolutionary histories.

In this study, we investigated whether ancient allopolyploidization has indeed contributed to the forming and establishment of major plant groups and whether such events have influenced phylogenetic relationships among early-diverging angiosperms. To this end, we analyzed genomic data from early-diverging angiosperm lineages that have experienced at most a single lineage-specific WGD event. By combining gene-tree phylogenetics and synteny-based biased fractionation, we identified and phased the subgenomes of putative allopolyploids to construct subgenome-resolved phylogenies. Integrating hundreds of nuclear gene trees with this subgenomic information, we demonstrate that the Chloranthales WGD represents an ancient allopolyploidization event involving progenitors related to both eudicots and monocots, revealing a previously unrecognized reticulate origin underlying a major angiosperm lineage. Our study provides a systematic subgenome-resolved investigation of ancient allopolyploidy in early angiosperms and highlights the importance of incorporating polyploid histories into phylogenomic frameworks for resolving deep evolutionary relationships.

## Results and Discussion

### Genome selection, WGD history, and syntenic pattern

We selected seven species encompassing seven of the eight early divergent lineages of angiosperms: *Amborella trichopoda* (Amborellales) (48), *Nymphaea colorata* (Nymphaeales) (49), *Illicium verum* (Austrobaileyales) (50), *Chloranthus spicatus* (Chloranthales) (11), *Aristolochia fimbriata* (magnoliids) (32), *Acorus gramineus* (early-diverging monocots) (51), and *Buxus austro-yunnanensis* (early diverging eudicots) (52). Crucially, these genomes were selected because they have experienced at most one detected whole-genome duplication (WGD) since the divergence of the most recent common ancestor of angiosperms. The Ceratophyllales lineage was deliberately excluded to avoid the analytical complexities introduced by its three independent WGD rounds. Two gymnosperms: *Thuja plicata* (53) and *Cycas panzhihuensis* (54), were included as outgroups (Figure 1a and Supplementary Table S1). All nine genomes selected demonstrated high completeness, with BUSCO scores exceeding 85% (Figure 1b).

**Figure 1.**
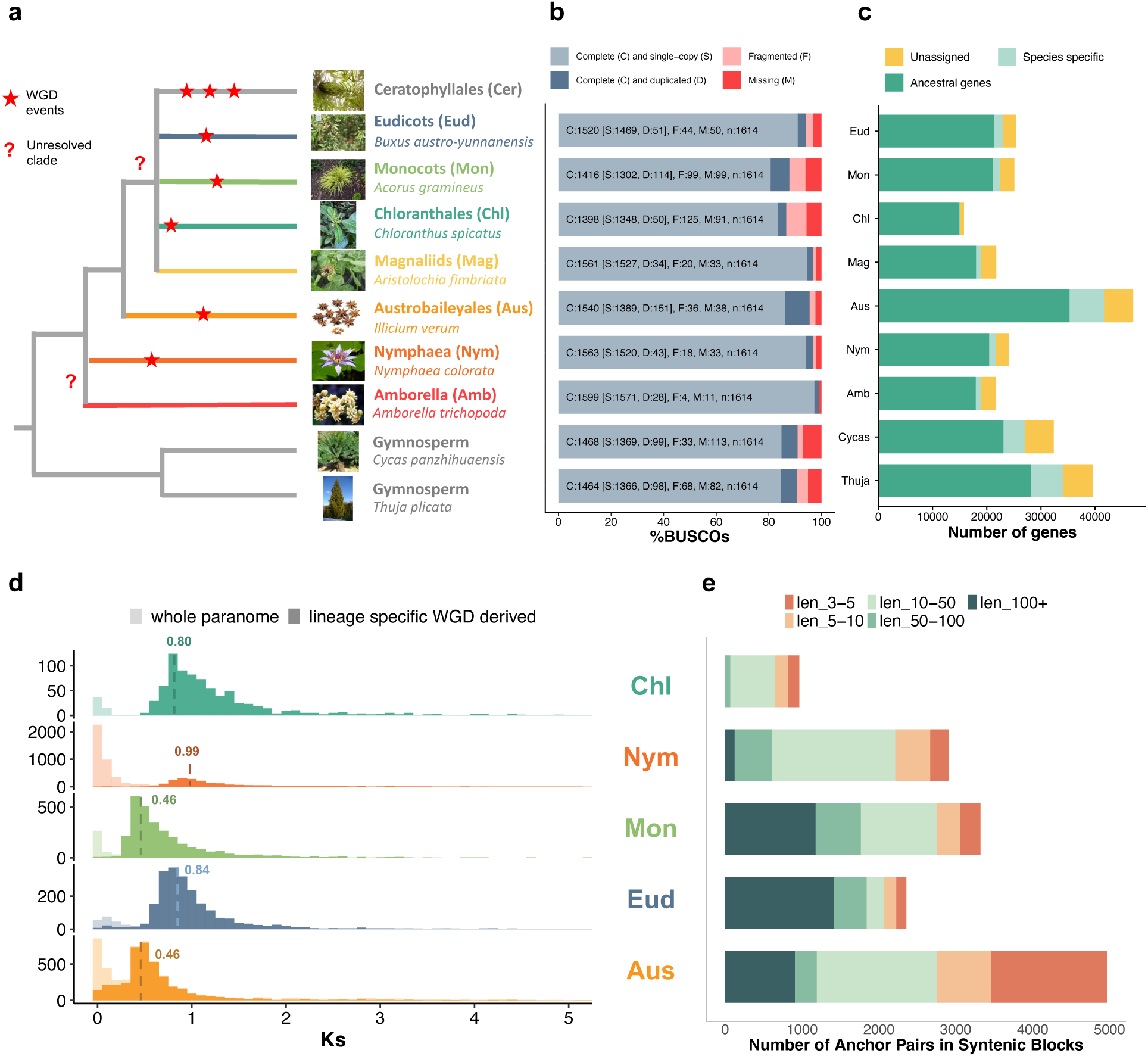
Phylogenetic relationships, genomic features and whole-genome duplication (WGD) history of major angiosperm lineages and selected representative taxa. **a)** A schematic phylogeny adapted from previous studies (3, 7, 12, 14). Seven genomes were selected to represent each major angiosperm lineage, excluding Ceratophyllales (which has experienced three rounds of WGD). Two gymnosperm genomes were included as outgroups. Red stars indicate WGD events that occurred after the divergence of the most recent common ancestor of angiosperms. Question marks denote topologically unresolved nodes reflecting conflicting results from prior studies. **b)** Genome assembly completeness of the nine selected genomes, assessed using Benchmarking Universal Single-Copy Orthologs (BUSCO; embryophyta_odb10) (55). **c)** Distribution of ancestral gene content across the nine genomes. Gene classification is based on orthogroup clustering. Dark green bars represent ancestral genes with homologs in at least one other sampled species; light green bars denote species-specific multi-copy genes (in-paralogs); yellow bars represent unassigned, species-specific single-copy genes. **d)** Synonymous substitution rate (*Ks*) distributions of lineage specific intragenomic syntenic paralogs (dark colors) and the whole paranome (light colors) to infer lineage-specific WGDs. Dashed lines and the numbers above them indicate *Ks* peaks identified via Gaussian mixture modeling using WGD v2 (--peak option) (56). **e)** Length distribution of syntenic blocks, represented by the proportion of blocks containing varying numbers of collinear anchor gene pairs, for the five genomes that have experienced lineage-specific WGDs subsequent to early angiosperm divergence.

Following our analytical workflow (Supplementary Figure S1), we performed orthogroup clustering across the nine genomes. Of the 252,868 total annotated genes, 88.3% were successfully clustered into 19,628 gene families, including 1,106 strict single-copy orthogroups shared across all taxa (Supplementary Table S2). To ensure the robustness of phylogenetic inference, gene families present in fewer than four taxa were filtered out, resulting in 13,569 gene families for subsequent gene tree construction. Concurrent with the clustering, we identified “ancestral genes”, defined as genes possessing homologs in at least one other lineage (Figure 1c, green stacked columns). The resulting gene trees and ancestral gene sets served as the fundamental dataset for subsequent analyses.

We then characterized the synteny patterns of the five genomes that underwent lineage-specific WGDs subsequent to angiosperm divergence. Intragenomic synteny analysis identified relatively well-preserved syntenic blocks in *I. verum*, *Ac. gramineus*, and *B. austro-yunnanensis*, exhibiting prominent synonymous substitution rate (*Ks*) peaks at 0.46, 0.46, and 0.84, respectively. In stark contrast, *C. spicatus* (*Ks* peak = 0.80) and *N. colorata* (*Ks* peak = 0.99) exhibited highly fragmented syntenic structures (Figure 1d; Supplementary Figure S2). This fragmentation was further corroborated by quantitative assessments: the proportion of syntenic genes in *C. spicatus* (12%) and *N. colorata* (25%) was generally lower, and their syntenic blocks were markedly shorter than those in the other genomes (19%–27%) (Figure 1e; Supplementary Figure S3). These metrics suggest that the *C. spicatus* and *N. colorata* lineages have experienced extensive post-WGD chromosomal rearrangements.

### Phylogenetic evidence reveals an allopolyploid origin for the Chloranthales WGD

To test if the five focal lineages have an allopolyploid or autopolyploid ancestry, we employed a phylogenetic framework based on the tree topology of anchor gene pairs. Here, we define “anchor gene pairs” as pairs of duplicated genes located on corresponding segments within a syntenic block. To ensure robust phylogenetic inference, we first excluded a subset of anchor pairs (shown in grey as "Filtered" in Figure 2) that met any of the following criteria: (1) the two genes did not belong to the same orthogroup across the analyzed genomes; (2) an anchor gene appeared more than once across different syntenic blocks; or (3) the anchor pair belonged to an orthogroup containing fewer than four taxa, thereby precluding reliable gene tree reconstruction.

**Figure 2.**
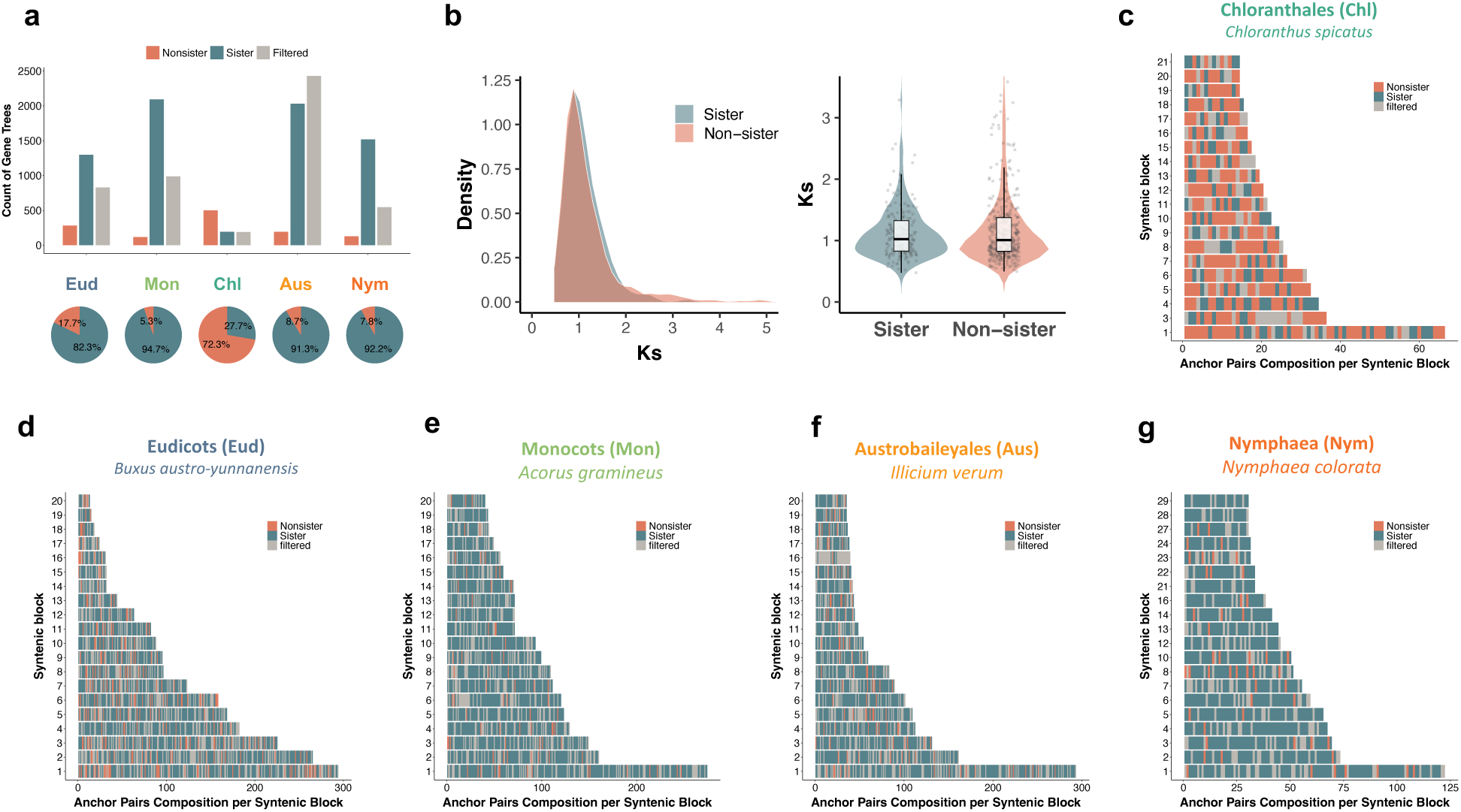
Genomic evidence for allopolyploid origin of Chloranthales. **a)** Counts and proportions of WGD-derived anchor gene pairs exhibiting sister versus non-sister topologies across the five lineages with lineage-specific WGDs. The three topological categories are defined as follows. Sister: anchor gene pairs that cluster as sister taxa to each other before clustering with orthologs from other lineages. Non-sister: anchor gene pairs that cluster with orthologs from other lineages prior to forming a sister relationship with each other. Filtered: anchor gene pairs excluded from the analysis because (1) the two duplicate genes did not cluster into the same orthogroup, (2) an anchor gene mapped to multiple different syntenic blocks, or (3) the pair belonged to an orthogroup containing fewer than four taxa, thereby precluding reliable gene tree reconstruction. Abbreviations: Eud, early-diverging eudicots; Mon, early-diverging monocots; Chl, Chloranthales; Aus, Austrobaileyales; Nym, Nymphaeales. **b)** Comparison of *Ks* distributions between sister- and non-sister-grouped anchor pairs in *Chloranthus spicatus.* **c-g)** Genomic distribution of the three anchor gene pair categories across the 20 longest syntenic blocks in **c)** *Chloranthus spicatus* (Chloranthales), **d)** *Buxus austro-yunnanensis* (eudicots), **e)** *Acorus gramineus* (Basal monocots), **f)** *Illicium verum* (Austrobaileyales) and **g)** *Nymphaea colorata* (Nymphaeales).

Focusing solely on the successfully retained (unfiltered) anchor pairs, we quantified the proportion of these WGD-derived anchor pairs that clustered as sister taxa versus non-sister taxa, using the gene family trees constructed across all nine genomes. In four of the lineages, i.e., monocots (94.7% of retained pairs), eudicots (82.3%), Austrobaileyales (91.4%), and Nymphaeales (92.2%), the vast majority of anchor pairs clustered as sisters. In contrast, Chloranthales exhibited the opposite pattern: among its retained anchor pairs, only 27.7% formed sister clades, while 72.3% formed non-sister group relationships (Figure 2a). Because autopolyploidy typically yields sister relationships (reflecting duplication within the same genome), whereas allopolyploidy results in non-sister clustering (reflecting divergence between different parental subgenomes), these topological signals indicate that the WGD in Chloranthales should be an ancient allopolyploid event. Conversely, the lineage-specific WGDs observed in the other four lineages exhibit patterns consistent with autopolyploidy. Although we cannot exclude the possibility that this autopolyploid-like signal is an artifact of limited taxonomic sampling, which might miss extinct or unsampled diploid progenitors, this interpretation is supported by independent evidence. Specifically, analysis of gene-tree discordance by Huang et al. (2025) (9) supported the monophyly of eudicots, monocots, and magnoliids, consistent with our finding that the WGDs in the other four early-diverging lineages likely represent autopolyploidization events.

To exclude the possibility that the observed alloploidization pattern of Chloranthales was artifactual and to clarify the factors underlying the frequent non-sister clustering of anchor pairs, we systematically evaluated potential methodological and interpretative biases. First, because the *Ks* distribution of Chloranthales is broader than that of other taxa, we tested whether substitution rate variation could account for the elevated proportion of non-sister groupings. Comparison of *Ks* distributions between sister- and non-sister-grouped anchor pairs revealed complete overlap, indicating that rate heterogeneity does not explain the unexpectedly high frequency of non-sister clustering in Chloranthales (Figure 2b). Second, we tested the effect of alternative taxon sampling within Chloranthales. Repeating orthogroup inference, syntenic block detection, and gene tree reconstruction using *Chloranthus sessilifolius* (12) instead of *Chloranthus spicatus* yielded a similar pattern of sister relationships among anchor gene pairs (Supplementary Figure S4a). Third, we examined the robustness of our results to model choice in gene tree reconstruction. Instead of applying the JTT+R model, we used the ModelFinder (MFP) approach in IQ-TREE2 to select the best-fitting substitution model for each gene family. The sister grouping pattern of anchor pairs from resulting topologies were highly consistent with those from topologies inferred under JTT+R (Supplementary Figure S4b).

Next, we investigated whether the profound non-sister clustering in Chloranthales could be driven by other complex biological factors, primarily differential gene loss (hidden paralogy) and incomplete lineage sorting (ILS). We first assessed the potential confounding effect of hidden paralogy stemming from older WGD events. Currently, there is no evidence of a WGD event uniquely shared by mesangiosperms. Rather, it is well established that all angiosperms share at least one ancient WGD with gymnosperms, although whether a subsequent angiosperm-specific WGD occurred remains debated (27–31). Attributing the observed non-sister pattern to ancient hidden paralogy requires the genome-wide persistence of numerous ancient paralogs—a highly improbable evolutionary scenario. This would necessitate both the widespread biased loss of duplicates from the recent Chloranthales-specific WGD and the intact retention of ancestral WGD paralog pairs through subsequent duplication events to the present day. There is no known biological mechanism to systematically maintain such a pattern genome-wide. More importantly, paralogs derived from an ancient, pre-angiosperm WGD would exhibit significantly higher *Ks* values compared to those derived from the Chloranthales-specific WGD. As shown in Figure 2b, the *Ks* distributions for non-sister and sister anchor paralogs overlap entirely, confirming a synchronous origin and refuting the hidden paralogy hypothesis.

We next investigated whether incomplete lineage sorting (ILS) following an autopolyploidization event could explain the non-sister clustering of anchor gene pairs in Chloranthales. Under the multi-species coalescent model, the expected ratio of sister (concordant) to non-sister (discordant) topologies among duplicates depends on the number of radiating lineages. For a three-taxon system, a pure ILS model predicts a minimum of one-third (33.3%) sister and a maximum of two-thirds (66.7%) non-sister relationships. If the Chloranthales WGD occurred prior to a radiation of four or more lineages, the expected sister ratio could drop to 25% or lower. However, interspecific *Ks* profiles show that the divergence peaks of Chloranthales with eudicots and monocots are close to the intraspecific WGD peak of Chloranthales, whereas the peak with magnoliids is older (Supplementary Figure S5). This suggests that any post-WGD ILS process likely involved no more than three lineages, supporting the relevance of the 1:2 theoretical threshold. Because our observed ratio (27.7% sister vs. 72.3% non-sister) falls below this 33.3% minimum, ILS under a three-taxon model appears insufficient to explain the pattern. Therefore, while ILS likely contributed to gene tree discordance, it is unlikely to be the sole driver, pointing instead toward an ancient allopolyploid origin.

In principle, a true allopolyploidization event would initially yield zero WGD-derived sister pairs. However, homoeologous exchanges (HEs) and gene conversion, wherein a dominant homoeologous chromosome replaces its recessive counterpart, can subsequently obscure subgenome-specific signals, causing some allopolyploid-derived anchor pairs to cluster as sisters. Taken together, these syntenic and phylogenetic lines of evidence demonstrate that the WGD in Chloranthales represents a definitive case of ancient allopolyploidization, followed by extensive HEs that partially homogenized the two distinct parental subgenomes. Moreover, we demonstrated that taxon sampling, substitution model choice, among-gene substitution rate variation, ILS, and hidden paralogy do not account for the observed allopolyploidization pattern in Chloranthales.

In the other four lineages, we observed the similar mosaic pattern, albeit with the vast majority of anchor gene pairs clustering as sisters (Figures 2d-g), this spatial homogeneity aligns with the high proportions of sister-grouping anchor pairs quantified earlier (Figure 2a) and reinforces the interpretation of less reticulate, putatively autopolyploid origins for these specific WGDs. Nevertheless, we did observe a minor fraction of discordant anchor pairs, representing 648 non-redundant gene trees in total, that did not cluster as sister groups within these putatively autopolyploid lineages (specifically, 118 trees in *Ac. gramineus*, 280 in *B. austro-yunnanensis*, 191 in *I. verum*, and 128 in *N. colorata*). As discussed above, this topological discordance could be driven by several confounding factors, including substitution rate variation between paralogs, ILS, hidden paralogy, or gene tree estimation errors. To ensure the highest stringency, we completely removed the 648 gene trees containing these non-sister anchor pairs from other four lineages prior to downstream analyses.

### Sub-genome phasing of Chloranthales based on biased fractionation

Subgenome dominance is a widespread phenomenon in polyploids, particularly in allopolyploids, characterized by asymmetric evolutionary trajectories between parental subgenomes (57–59). Typically, one subgenome undergoes preferential gene loss (fractionation), whereas the other retains a greater proportion of ancestral genes and is therefore considered dominant. Furthermore, genes retained on the more fractionated subgenome often exhibit reduced expression levels relative to their homoeologous counterparts on the dominant subgenome (57, 60).

To characterize the evolutionary trajectory of the allopolyploidization event in Chloranthales, we leveraged patterns of subgenome-biased fractionation to phase the genes originating from the two parental subgenomes. Theoretically, each WGD-derived syntenic block comprises two homoeologous segments. Within each block, the segment retaining a higher number of unique ancestral genes (previously defined as having homologs in at least one other sampled lineage) was designated as the dominant subgenome, whereas the segment with fewer unique ancestral genes was classified as the recessive subgenome. To rigorously assign subgenome identities, we applied a Chi-square (χ^2^) test to evaluate whether the disparity in ancestral gene counts between paired segments was statistically significant (*p*<0.05) (Figure 3a; see also Methods and Supplementary Figure S6).

**Figure 3.**
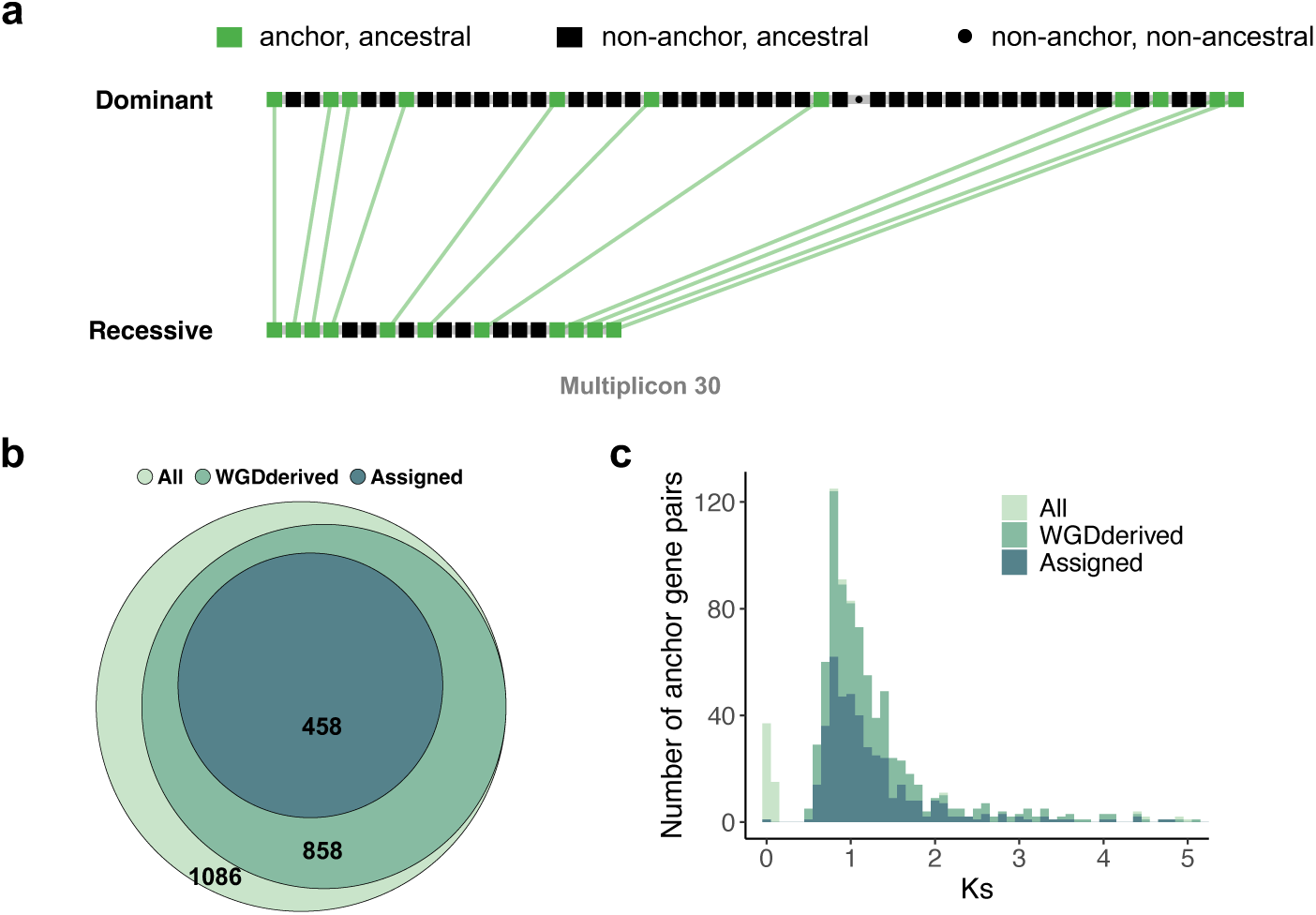
Subgenome phasing based on biased fractionation following lineage-specific WGD of Chloranthales. **a)** Schematic illustration of dominant and recessive subgenomes assigned based on biased gene retention (fractionation), using one representative syntenic block (Multiplicon 30) from *Chloranthus spicatus* as an example. **b)** Venn diagram illustrating the nested subsets of anchor gene pairs assigned to subgenomes in *Chloranthus spicatus*. **c)** Synonymous substitution rate (*Ks*) distributions of different categories of anchor gene pairs before and after subgenome phasing, demonstrating no distinct age bias. The plotted categories include: All, the total number of syntenic anchor pairs identified in the *C. spicatus* genome; WGD-derived, the subset of anchor pairs specifically derived from the lineage-specific WGD event; and Assigned, the final curated subset of WGD-derived anchor pairs successfully phased into dominant and recessive subgenomes based on biased fractionation.

We observed extensive gene fractionation and genomic rearrangement following the Chloranthales WGD, as indicated by the predominance of non-anchor genes (singlets) relative to retained anchor genes (collinear homoeologous pairs) across WGD-derived regions. Following this statistical framework, among 858 WGD-derived gene pairs distributed across 89 syntenic blocks, 458 gene pairs from 27 syntenic blocks were successfully assigned to their respective subgenomes (Figure 3b; Supplementary Table S3). Importantly, the *Ks* distributions of subgenome-assigned anchor gene pairs closely matched those of all WGD-derived anchor gene pairs, suggesting that our assignment procedure did not preferentially capture duplicates from specific evolutionary age classes (Figure 3c). These curated subgenome-resolved gene sets were subsequently used for downstream analyses.

### Ancestral eudicot and monocot lineages as putative parental contributors to Chloranthales allopolyploidization

To infer the putative progenitor lineages involved in the Chloranthales allopolyploidization event, we focused on gene trees containing paralogous gene pairs generated by the Chloranthales-specific WGD. We filtered WGD-derived anchor gene pairs that (i) belonged to the same orthogroup and (ii) retained orthologs in at least four focal taxa. Based on these filtered gene pairs, we generated three datasets according to their phylogenetic configurations: 311 gene trees exhibiting non-sister relationships between Chloranthales paralogs (Dataset i), 154 gene trees exhibiting sister relationships (Dataset ii), and 165 gene trees in which Chloranthales paralogs could be assigned to different subgenomes (Dataset iii).

We first performed a statistical assessment of gene-tree topologies to identify the most strongly supported sister lineage for each Chloranthales paralog using Dataset i. Among the 311 non-sister gene trees containing all four mesangiosperm taxa, in which at least one Chloranthales paralog was resolved as sister to a non-Chloranthales lineage, eudicots and monocots emerged as the two most frequently supported sister groups. The support for these two relationships was nearly balanced, with 111 trees supporting a sister relationship with monocots and 109 supporting a sister relationship with eudicots (Figure 4a). Partitioning these gene trees according to subgenome identity provided clearer resolution: Across the 165 gene trees (Dataset iii), the dominant subgenome was more frequently associated with eudicots, whereas the recessive subgenome was more frequently associated with monocots (Figure 4b). To further evaluate this parental assignment, we examined the 154 gene trees in Dataset ii, where the two Chloranthales paralogs formed sister relationships. Such topologies may arise from homoeologous exchange (HE), during which gene conversion or reciprocal exchange between homoeologous chromosomes can alter the genomic representation of parental subgenomes 39. Under this scenario, HE-derived gene pairs are expected to retain stronger phylogenetic affinity with the dominant parental lineage (eudicots). Consistent with this expectation, the putative dominant parent (eudicots) was recovered as the sister lineage to Chloranthales in the majority of these gene trees (Supplementary Figure S7). Together, these complementary tree-based analyses provide strong evidence that eudicots and monocots represent the most plausible parental lineages contributing to the ancient Chloranthales allopolyploidization event (Figure 4c).

**Figure 4.**
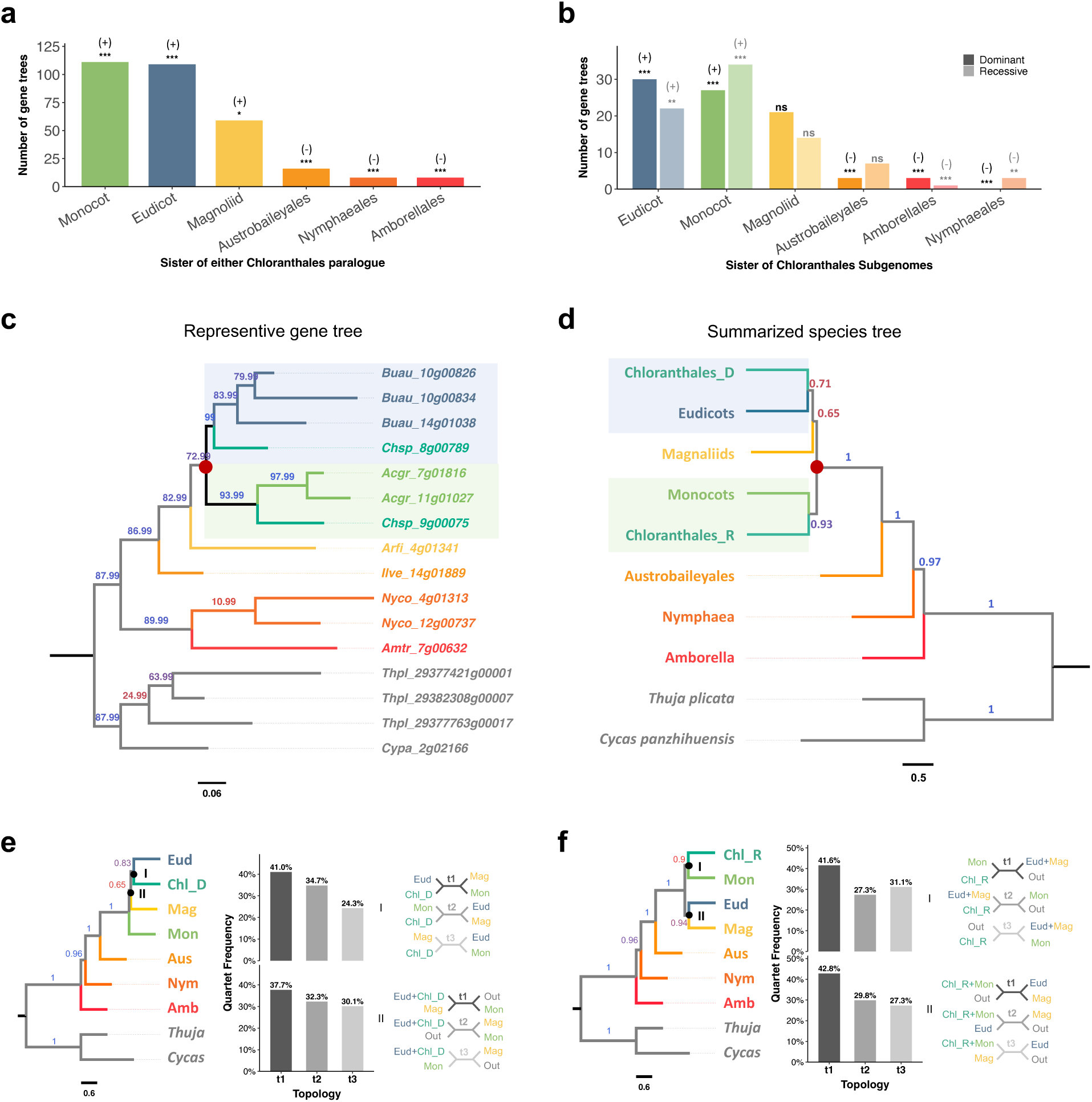
Assessment of parental lineages of the Chloranthales allopolyploidization. **a)** Statistical evaluation of the closest sister lineages inferred from 311 non-sister gene trees containing all four mesangiosperm taxa. The y-axis represents the absolute count of gene trees supporting each respective lineage (x-axis) as the sister group. Signals (+/-) above each bar indicate observed frequencies that are significantly higher (+) or lower (-) than expected by chance, respectively. Significance markers denote Benjamini-Hochberg FDR-corrected *P*-values from binomial tests (\*\*\**P*_adj_ *<0.001*; \*\**P*_adj_ *<0.01*; \**P*_adj_ < 0.05; ns, not significant). **b)** Statistical evaluation of the closest sister lineages inferred from 165 subgenome-phased gene trees. The y-axis shows the number of gene trees supporting the sister relationship for paralogs from either the dominant subgenome (dark shading) or the recessive subgenome (light shading) of Chloranthales. Significance markers are defined as in **a)**. **c)** A representative gene tree topology illustrating the allopolyploid origin of Chloranthales (OG0001793_HOG0004998). The red node indicates the divergence of the two putative parental lineages (eudicots and monocots); the hybridization/WGD event occurred subsequent to this divergence. Node labels represent ultrafast bootstrap (UFBoot) support values. Scale bar indicates substitutions per site. **d)** Subgenome-resolved species tree inferred from 182 gene trees using WASTRAL. To maximize gene tree completeness, a single anchor gene per orthogroup was randomly selected for the other four lineages, regardless of subgenomic origin. The red circle denotes the divergence of the two putative parental lineages. Branch labels represent local posterior probabilities. Tip branch lengths are arbitrary and not phylogenetically meaningful (65). **e)** Quartet frequencies of the three alternative topologies at two nodes (I and II) within mesangiosperms, calculated across the 165 subgenome-phased gene trees while retaining only the dominant subgenome of Chloranthales. **f)** Quartet frequencies of the three alternative topologies at two nodes (I and II) within mesangiosperms, calculated across the 165 subgenome-phased gene trees while retaining only the recessive subgenome of Chloranthales.

Next, based on the 165 non-sister gene trees with resolved Chloranthales subgenome identities (Dataset iii), we reconstructed a multilabeled species tree of early-diverging angiosperms at the subgenome level. The species tree inferred using wASTRAL recovered the same pattern, with the dominant Chloranthales subgenome clustering with eudicots and the recessive subgenome clustering with monocots (Figure 4d). In contrast, the species tree inferred from unphased multi-copy gene trees placed Chloranthales as sister to eudicots (Supplementary Figure S8), consistent with previous phylogenomic analyses (17, 61). This discrepancy highlights the improved phylogenetic resolution gained from subgenome-aware approaches, which are critical for resolving deep evolutionary relationships affected by reticulate evolution.

Given that several mesangiosperm nodes exhibited low local posterior support and short internal branches, we investigated whether incomplete lineage sorting (ILS) contributed to the observed gene-tree discordance. We analyzed datasets containing only the dominant (Figure 4e) or recessive (Figure 4f) Chloranthales subgenome. Under the multispecies coalescent model, if gene-tree discordance is driven solely by ILS, the two alternative discordant topologies for a given species triplet are expected to occur at approximately equal frequencies (62). For the relationships among eudicots, monocots, and magnoliids, the two discordant topologies occurred at nearly equal frequencies, suggesting that ILS represents a major contributor to phylogenetic conflict among these mesangiosperm lineages. In contrast, for relationships involving the phased Chloranthales subgenomes, the two discordant topologies showed a similar but not perfectly equal frequency distribution, suggesting that although ILS likely explains a substantial proportion of the observed gene-tree discordance, additional evolutionary processes may also have contributed.

The phylogenetic position of Chloranthales has long remained unresolved. The APG III system positioned Chloranthales as the sister group to magnoliids (63), whereas APG IV demoted it to an unresolved polytomy alongside magnoliids and the monocot-eudicot-Ceratophyllales clade due to weak statistical support (3). Previous phylogenomic analyses have converged on three conflicting topologies: placing Chloranthales as sister to eudicots (17, 61), sister to magnoliids (8, 11, 12, 15, 64), or as the basalmost lineage of mesangiosperms (10, 13, 14, 16, 18). Notably, despite these shifting placements, excluding Chloranthales from these datasets consistently restores an identical, highly stable topology among the remaining mesangiosperm clades (eudicots, *Ceratophyllales*, magnoliids, and monocots) (11, 12, 17, 61, 64). While recent genomic analyses have implicated ancient hybridization as the primary driver of deep mesangiosperm discordance over ILS (9, 11, 12), the precise evolutionary nature of this reticulation has remained elusive. Our study contextualizes these previous findings by demonstrating that this confounding phylogenetic signal is actually the genomic signature of an ancient allopolyploidization involving eudicot- and monocot-related progenitors, thereby providing a mechanistic explanation for its historically unstable phylogenetic placement.

Our findings provide evidence that ancient hybridization can occur between deeply diverged lineages and leave detectable phylogenomic signatures over geological timescales. Hybridization and polyploidization are major evolutionary forces driving plant diversification, speciation, and adaptive radiation (21–25). Approximately 25% of plant species have been estimated to be involved in hybridization (21), and at least 15% of angiosperm speciation events are associated with polyploidy (22), although these estimates may represent conservative lower bounds. Despite the importance of allopolyploidization, which combines hybridization with whole-genome duplication, its contribution to deep-time phylogenomic evolution has historically received limited attention. This neglect likely stems from the assumption that ancient allopolyploid events are difficult to detect in deeply diverged lineages because their original genomic signals are progressively eroded through extensive fractionation, chromosomal rearrangement, and subgenome homogenization (41, 42). However, studies in other kingdoms, including landmark analyses of ancient yeast polyploidizations, have demonstrated that allopolyploid origins can be successfully reconstructed even after hundreds of millions of years of divergence (39). Our study provides the first systematic assessment of how ancient allopolyploidization can contribute to phylogenetic discordance among major angiosperm lineages. More broadly, these findings highlight the importance of incorporating reticulate evolutionary histories, particularly ancient allopolyploid events, into models of deep phylogenetic reconstruction across major angiosperm clades.

### Concluding remarks

Allopolyploidization, which combines genome duplication with hybridization, represents a major source of complexity for phylogenetic reconstruction, particularly in deeply diverged lineages. In this study, we demonstrate that the Chloranthales lineage originated through an ancient allopolyploidization event involving monocot- and eudicot-related ancestral lineages. This discovery suggests that allopolyploid speciation between deeply diverged clades may have played a more important and previously underappreciated role in shaping early angiosperm evolution. Although our analyses are constrained by current methodological limitations and available genomic resources, continued advances in ancient polyploid subgenome resolution, together with the expanding availability of high-quality genome assemblies from early-diverging lineages, will provide increasingly powerful opportunities to test, refine, and extend our understanding of ancient reticulate evolution in angiosperms.

## Materials and Methods

### Genome selection and quality assessment

We selected one representative genome from each of seven early-diverging angiosperm lineages, which have experienced no or only one WGD event since the divergence of the ancestor of angiosperms, excluding *Ceratophyllales*, which has undergone three rounds of whole-genome duplication (WGD) following its divergence from the angiosperm ancestor (66).

The selected genomes are from *Amborella trichopoda* (Amborellales) (48), *Nymphaea colorata* (Nymphaeales) (49), *Illicium verum* (Austrobaileyales) (50), *Chloranthus spicatus* (Chloranthales) (11), *Aristolochia fimbriata* (magnoliids) (32), *Acorus gramineus* (early-diverging monocots) (51), and *Buxus austro-yunnanensis* (early-diverging eudicots) (52). Additionally, we selected two outgroup species from gymnosperms: *Thuja plicata* (53) and *Cycas panzhihuensis* (54). Accession numbers and references for all genomes are provided in Supplementary Table S1. Genome assembly completeness was re-evaluated using BUSCO (embryophyta_odb10) (55).

### Orthogroup clustering and ancestral gene identification

Prior to orthogroup clustering, the gene annotations for all nine genomes were pre-processed by retaining only the longest transcript (isoform) per gene locus and removing sequences that contained internal premature stop codons. Orthogroup inference was performed using OrthoFinder v3.0.1b1 with 36 threads (67). Based on the hierarchical orthogroup clustering results (N0.tsv), we defined ’ancestral genes’ as those exhibiting homology with at least one other species. Therefore, any gene assigned to an orthogroup containing sequences from multiple species was classified as ancestral for that given lineage.

### Intragenomic synteny analysis and paleologs identifying

Both intergenomic and intragenomic synteny blocks were identified across and within the nine genomes using the WGD v2 pipeline (56). First, the “*wgd dmd*” module was executed to define gene families using the Markov Cluster Algorithm (MCL) with an inflation factor of 3 (-I 3). Subsequently, collinearity detection was performed using the “wgd syn” module, which internally invokes I-ADHoRe v3 (68) to identify homologous genomic regions based on conserved gene content and order (terms used interchangeably with ’synteny’ in this study). To specifically capture duplications originating from whole-genome duplications, we restricted our analysis to level-2 multiplicons (i.e., syntenic regions retaining exactly two paralogous copies). The precise parameters passed to I-ADHoRe were: --iadhore_options level_2_only=true, gap_size=40, cluster_gap=40, q_value=0.70. The minimal number of anchor points in a cluster was set to three as default.

To isolate paleologs, specifically, duplicated genes retained from the most recent ancient polyploidy event (hereafter referred to as the lineage-specific WGD), we combined synteny networks with paralog age distributions. The synonymous substitution rate (*Ks*) for all paralogous pairs was calculated using the “wgd ksd” module under default settings. Lineage-specific WGD peaks were then statistically delineated via Gaussian mixture modeling using the wgd peak module, employing a node-weighted approach for redundancy resolution (--weighted). Paleologs associated with these WGD events were robustly extracted based on the 95% confidence interval (CI) of the mean *Ks* calculated across all syntenic gene pairs anchored within a given segment (the SegmentGuideKs metric in WGD v2). These filtered paleologs were utilized for downstream subgenome phasing.

Phylogenetic assessment of anchor gene pair topologies

To determine whether the lineage-specific WGDs in the five focal genomes originated from allo-or autopolyploidy, we employed a phylogenetic framework to assess the topological relationships within each anchor gene pair. First, phylogenetic trees were reconstructed for all identified orthogroups except those with fewer than four taxa, resulting in 13,569 gene families. Protein sequences were aligned using MAFFT v7.525 (69), with the L-INS-i or FFT-NS-i strategy automatically selected (--auto), and the resulting alignments were trimmed to remove spurious regions using TAPER v1.0.2 under default settings (70). Maximum likelihood (ML) phylogenies were inferred using IQ-TREE v2.3.6 (71). To evaluate the robustness of our results to model selection, we ran two parallel tree-building analyses: one applying a uniform JTT+R model, and another utilizing ModelFinder (-m MFP) to automatically determine the best-fitting substitution model for each orthogroup. To optimize tree topology, a more thorough NNI search (-bnni) was applied. Branch support was evaluated using 1,000 SH-aLRT replicates and 1,000 ultrafast bootstrap replicates (-alrt 1000 -bb 1000).

To ensure the accuracy of topological interpretations, we rigorously filtered the anchor pairs prior to analysis. Pairs were excluded if they met any of the following criteria: (1) the two duplicate genes did not cluster into the same orthogroup; (2) either gene within the pair mapped to multiple different syntenic blocks; or (3) the anchor pair was assigned to an orthogroup containing fewer than four taxa, which precludes meaningful phylogenetic reconstruction. Using custom Python scripts, the successfully retained WGD-derived anchor pairs for each species were classified based on their tree topologies, namely, whether the two paralogs formed a monophyletic sister clade before clustering with orthologs from other lineages. Finally, the overall proportions of sister versus non-sister topologies were quantified using Python, and the spatial distribution of these topological categories across genomic syntenic blocks was visualized using custom R scripts.

### Subgenome phasing of Chloranthales based on biased fractionation

Building upon the intragenomic synteny networks and orthogroup clustering established earlier, we phased the anchor gene pairs derived from the Chloranthales-specific WGD into distinct subgenomes. Building upon the intragenomic synteny networks and orthogroup clustering established earlier, we pahsed the anchor gene pairs derived from the Chloranthales-specific WGD into distinct subgenomes. This phasing procedure relies on the differential retention of genes (biased fractionation) and was adapted from well-established methodological frameworks (72, 73) (Supplementary Figure S6).

Theoretically, each WGD-derived syntenic block consists of two homoeologous segments. For each segment, we partitioned the genomic space into anchor (collinear pairs) and non-anchor (singlets) regions, and subsequently classified the genes within these regions as either ancestral or non-ancestral. To phase the subgenomes, we specifically quantified the number of ancestral genes (defined as uniquely retained genes possessing orthologs in at least one of the other eight sampled genomes) situated within the non-anchor regions of each segment (depicted as black circles in Supplementary Figure S6). Within each paired syntenic block, the segment retaining a numerically superior set of these non-anchor ancestral genes was designated as the dominant subgenome, while its homoeologous counterpart with fewer retained genes was classified as the recessive subgenome. The statistical significance of the disparity in gene retention (used as a proxy for segment length) between the two corresponding segments was rigorously assessed using a Chi-square (χ2) test (*p*<0.05). Only after this subgenome phasing was statistically validated were the syntenic anchor genes (depicted as green circles in Supplementary Figure S6) confidently extracted and assigned to their respective subgenomes for subsequent downstream analyses.

### Identification of the putative parental lineages of the Chloranthales allopolyploid

To infer the putative parental lineages, or at least close relatives, of the Chloranthales allopolyploid while minimizing the confounding effects of homoeologous exchanges (HEs) and gene conversion, we focused exclusively on gene trees containing WGD-derived Chloranthales paralogs with non-sister topologies. We applied additional strict filtering criteria to exclude: (1) large gene trees in which any of the nine genomes contained more than 10 homologous genes, and (2) trees where the homoeologs of any of the other four polyploid genomes (excluding Chloranthales) failed to cluster as sister groups. After this filtering pipeline, 421 gene trees were retained. Within this dataset, the Chloranthales paralogs in 251 gene trees were successfully phased into subgenomes based on biased fractionation.

We subsequently constructed three datasets for downstream analysis: (i) the 421 non-sister gene trees containing Chloranthales-specific WGD-derived homoeologs, (ii) the 251 non-sister gene trees containing subgenome-assigned homoeologs, and (iii) the 192 sister gene trees containing Chloranthales-specific WGD-derived homoeologs. Species labels were appended to the terminal branches of all trees, and the two Chloranthales anchor genes were assigned distinct labels to differentiate the paralogs. We used DISCO (Willson et al., 2022) under default settings to extract single-copy gene trees from the multi-copy gene families. Because our primary objective was to resolve the parental lineages within mesangiosperms, we restricted downstream analyses to trees that contained all four focal mesangiosperm clades (represented by five taxa, treating the two phased Chloranthales subgenomes as separate lineages): *Aristolochia fimbriata* (magnoliids), *Acorus gramineus* (monocots), *Buxus austro-yunnanensis* (eudicots), and the two subgenomes of *Chloranthus spicatus* (Chloranthales). Outgroups consisting of ANA-grade lineages and two gymnosperms were also required to be present. This rigorous taxon-filtering step yielded a final set of 311 non-sister single-copy trees with WGD-derived homoeologs (Dataset i), 154 sister single-copy trees with WGD-derived homoeologs (Dataset ii), and 165 non-sister single-copy trees with subgenome-assigned homoeologs (Dataset iii).

Using these three datasets, we performed statistical topological assessments of the gene trees to identify the most strongly supported sister lineage for each of the two Chloranthales anchor genes. For Dataset (i), we recorded all resolved sister lineages for either of the two anchor paralogs. For Dataset (ii), we recorded all sister lineages for the common ancestral node of two Chloranthales anchor genes. For Dataset (iii), sister lineages were resolved separately for anchor genes assigned to the dominant (D) or recessive (R) subgenomes. To identify sister lineages occurring significantly more or less frequently than expected by chance, a two-sided binomial test was conducted for each dataset. Multiple testing was corrected using the Benjamini-Hochberg false discovery rate (FDR) algorithm with the significance threshold set at alpha = 0.05 (adjusted *P*-value).

Second, using the 165 subgenome-assigned single-copy trees with full mesangiosperm coverage, we reconstructed a multispecies coalescent species tree. We employed the two-step weighted ASTRAL (wASTRAL v1.22.3.7) model (65) with default hybrid weighting (--mode 1), specifying support value thresholds of maximum (-x 100) and minimum (-n 0). Because wASTRAL does not estimate terminal branch lengths, we subsequently used ASTRAL-IV (74) under default settings to infer phylogenetically meaningful branch lengths based on the exact topology returned by wASTRAL. For comparison, we also used ASTRAL-Pro (74, 75) to infer a species tree directly from the unphased, multi-copy gene trees, allowing us to contrast the phylogenetic resolution and topological signals between the phased and unphased datasets.

## Supplementary Information

**Supplementary Figure S1.** Workflow of data analyses in this study.

**Supplementary Figure S2.** Intragenomic synteny dot plots illustrating whole-genome duplication (WGD) patterns in representative early-diverging angiosperms. (a) *Illicium verum* (Austrobaileyales), (b) *Acorus gramineus* (Basal monocots), (c) *Buxus austro-yunnanensis* (Basal eudicots), (d) *Nymphaea colorata* (Nymphaeales), (e) *Chloranthus spicatus* (Chloranthales) and (f) *Chloranthus sessilifolius* (Chloranthales). The Oxford grids display self-synteny based on gene rank, with synonymous substitution rate (*Ks*) values of syntenic paralogs color-coded according to the gradient legend.

**Supplementary Figure S3.** Proportion of syntenic blocks for five genomes representative for lineages with additional WGD.

**Supplementary Figure S4.** Genomic distribution of anchor gene pair categories on 20 longest syntenic blocks of **(a)** *Chloranthus sessilifolius* (Chloranthales), and **(b)** *C. spicatus* (Chloranthales) based on gene tree built with model MFP in IQTREE2. Three categories are defined as follows: Sister: anchor gene pairs that clustered as sisters to each other before clustering with orthologs from other lineages; Non-sister: anchor gene pairs that clustered with orthologs from other lineages prior to forming a sister relationship with each other; Filtered: anchor gene pairs excluded because they do not belong to the same orthogroup across the analyzed genomes, or the gene of anchor pair appeared more than once across different syntenic blocks, or anchor gene pairs within orthogroups containing fewer than four taxa, precluding gene tree reconstruction.

**Supplementary Figure S5. Interspecific *Ks* distributions between *Chloranthus spicatus* and representative mesangiosperm genomes.** The divergence *Ks* peaks of *C. spicatus* with eudicots (Chsp_Buau) and monocots (Chsp_Acgr) are closer to the intraspecific *Ks* peak of the *Chloranthus* WGD (Chsp_Chsp) than is the divergence *Ks* peak between *C. spicatus* and magnoliids (Chsp_Arfi). Abbreviations: Chsp, *Chloranthus spicatus*; Buau, *Buxus austro-yunnanensis*; Acgr, *Acorus gramineus*; Arfi, *Aristolochia fimbriata*.

**Supplementary Figure S6.** Subgenome phasing based on biased fractionation.

To enhance statistical power, the number of ancestral genes retained in non-anchor regions (black circles) was utilized as a proxy for the length (i.e., gene content) of each segment.

Significant differences in segment lengths between corresponding regions within each syntenic block were evaluated using a Chi-square (*χ*2) test (*p*<0.05). Following successful subgenome phasing, the anchor genes (green circles) were extracted for downstream analysis.

**Supplementary Figure S7.** Phylogenomic gene tree analysis supports Eudicots as the primary sister group to the Chloranthales clade, based on 154 sister grouping gene trees.

Distribution of sister group relationships for the Chloranthales clade derived from single-copy gene trees. The y-axis shows the absolute count of gene trees supporting each respective lineage (x-axis) as the sister group. Eudicots are recovered as the most frequent sister clade. Significance markers above each bar represent the results of binomial tests with Benjamini- Hochberg FDR correction (*** adjusted *P* < 0.05; ns, not significant).

**Supplementary Figure S8.** Species tree inferred by ASTRAL-Pro from multi-copy gene trees without subgenome phasing. The tree was reconstructed without phasing the subgenomes of the five polyploid species. Branch labels indicate bootstrap support values. Node labels represent the IDs of ancestral nodes (e.g., N0 denotes the most recent common ancestor (MRCA) of all nine lineages, whereas N2 denotes the MRCA of the seven angiosperm lineages).

**Supplementary Table S2.** Percentage and number of genes classified in different groups for each species

**Supplementary Table S2.** Percentage and number of genes classified in different groups for each species

**Supplementary Table S3.** χ2 test in subgenome assignment for each of the WGD-derived syntenic blocks in *Chloranthus spicatus*

## Acknowledgements

D.-Y.Z. acknowledges funding from the National Natural Science Foundation of China (32170223 and 31421063), the “111” Program of Introducing Talents of Discipline to Universities (B13008), Beijing Advanced Innovation Program for Land Surface Processes, and the National Key R&D Program of China (2017YFA0605104). Y.V.D.P. acknowledges funding from Ghent University (Methusalem funding, BOF.MET.2021.0005.01). Z.L. acknowledges funding from the Research Project of FWO (G0ADO25N) and the Special Research Grant from Ghent University (BOF.BAF.2024.0889.01). H.-C.C. acknowledges funding from the Research Foundation—Flanders (FWO) (No. 3G032219) and funding from the Walter Benjamin Programme of the German Research Foundation (DFG, No. 566819120, CH 3905/1-1). Y.C. acknowledges funding from China Scholarship Council (No. 202106040114).

## Author contributions

D.-Y.Z. and Y.V.D.P. conceived the research, Z.L. polished it. All analyses were performed by Y.C. and H.-C.C., Y.C. wrote the manuscript. D.-Y.Z., Y.V.D.P., Z.L. and H.-C.C. modified the manuscript. D.-Y.Z., Y.V.D.P. and Z.L. supervised the project.

## Declaration of generative AI in scientific writing

During the preparation of this manuscript, the authors used ChatGPT to assist with language editing. The authors subsequently reviewed and revised all AI-assisted text as necessary and take full responsibility for the final content of the published article.

## References

1. M. J. M. Christenhusz, J. Byng, The number of known plants species in the world and its annual increase. Phytotaxa 261, 201–217 (2016).

2. R. Govaerts, E. Nic Lughadha, N. Black, R. Turner, A. Paton, The World Checklist of Vascular Plants, a continuously updated resource for exploring global plant diversity. Sci. Data 8, 215 (2021).

3. G. The Angiosperm Phylogeny et al., An update of the Angiosperm Phylogeny Group classification for the orders and families of flowering plants: APG IV. Bot. J. Linn. Soc. 181, 1–20 (2016).

4. I. One Thousand Plant Transcriptomes, One thousand plant transcriptomes and the phylogenomics of green plants. Nature 574, 679–685 (2019).

5. S. Stefanović, D. W. Rice, J. D. Palmer, Long branch attraction, taxon sampling, and the earliest angiosperms: Amborella or monocots? BMC Evol. Biol. 4, 35 (2004).

6. T. Zhao et al., Whole-genome microsynteny-based phylogeny of angiosperms. Nat. Commun. 12, 3498 (2021).

7. G. Zhang, H. Ma, Nuclear phylogenomics of angiosperms and insights into their relationships and evolution. J. Integr. Plant Biol. 66, 546–578 (2024).

8. A. R. Zuntini et al., Phylogenomics and the rise of the angiosperms. Nature 629, 843–850 (2024).

9. X. Huang et al., Phylogenomic discordance is driven mainly by pervasive ancient hybridization and incomplete lineage sorting during the early divergence of major angiosperm lineages. Plant Commun. 6 (2025).

10. P. K. Endress, J. A. Doyle, Reconstructing the ancestral angiosperm flower and its initial specializations. Am. J. Bot. 96, 22–66 (2009).

11. X. Guo et al., Chloranthus genome provides insights into the early diversification of angiosperms. Nat. Commun. 12, 6930 (2021).

12. J. Ma et al., The *Chloranthus sessilifolius* genome provides insight into early diversification of angiosperms. Nat. Commun. 12, 6929 (2021).

13. H. Li et al., Plastid phylogenomic insights into relationships of all flowering plant families. BMC Biol. 19, 232 (2021).

14. H. Li et al., Origin of angiosperms and the puzzle of the Jurassic gap. Nat. Plants 5, 461–470 (2019).

15. M. J. Moore, C. D. Bell, P. S. Soltis, D. E. Soltis, Using plastid genome-scale data to resolve enigmatic relationships among basal angiosperms. Proc. Natl. Acad. Sci. U. S. A. 104, 19363–19368 (2007).

16. Y. Qiu et al., Angiosperm phylogeny inferred from sequences of four mitochondrial genes. J. Syst. Evol. 48, 391–425 (2010).

17. L. Yang et al., Phylogenomic insights into deep phylogeny of angiosperms based on broad nuclear gene sampling. Plant Commun. 1, 100027 (2020).

18. D. E. Soltis et al., Angiosperm phylogeny: 17 genes, 640 taxa. Am. J. Bot. **98**, 704–730 (2011).

19. J. H. Leebens-Mack et al., One thousand plant transcriptomes and the phylogenomics of green plants. Nature 574, 679–685 (2019).

20. M. D. Rasmussen, M. Kellis, Unified modeling of gene duplication, loss, and coalescence using a locus tree. Genome Res. 22, 755–765 (2012).

21. J. Mallet, Hybridization as an invasion of the genome. Trends Ecol. Evol. 20, 229–237 (2005).

22. T. E. Wood et al., The frequency of polyploid speciation in vascular plants. Proc. Natl. Acad. Sci. U. S. A. 106, 13875–13879 (2009).

23. G. W. Stull, K. K. Pham, P. S. Soltis, D. E. Soltis, Deep reticulation: the long legacy of hybridization in vascular plant evolution. Plant J. 114, 743–766 (2023).

24. D. C. Tank et al., Nested radiations and the pulse of angiosperm diversification: increased diversification rates often follow whole genome duplications. New Phytol. 207, 454–467 (2015).

25. S. Wu, Y. Wang, Z. Wang, N. Shrestha, J. Liu, Species divergence with gene flow and hybrid speciation on the Qinghai–Tibet Plateau. New Phytol. 234, 392–404 (2022).

26. P. S. Soltis, D. E. Soltis, Ancient WGD events as drivers of key innovations in angiosperms. Curr. Opin. Plant Biol. 30, 159–165 (2016).

27. Y. Jiao et al., Ancestral polyploidy in seed plants and angiosperms. Nature 473, 97–100 (2011).

28. C. Ruprecht et al., Revisiting ancestral polyploidy in plants. Sci. Adv. 3, e1603195.

29. A. Zwaenepoel, Y. Van de Peer, Inference of Ancient Whole-Genome Duplications and the Evolution of Gene Duplication and Loss Rates. Mol. Biol. Evol. 36, 1384–1404 (2019).

30. T. Shi, Y. Van de Peer, Revisiting ancient whole-genome duplications in the seed and flowering plants through the lens of dosage-sensitive genes. Sci. Adv. 12, eaea9797 (2026).

31. R. Zhang, M. A. Lysak, H.-Y. Shang, Y. Jiao, Y.-P. Ma, Orthologous synteny provides robust structural evidence for the ancestral angiosperm *ε*-WGD. bioRxiv [Preprint] (2026). http://biorxiv.org/content/early/2026/03/07/2026.03.05.709955.abstract ( (accessed 6 April 2026).

32. L. Qin et al., Insights into angiosperm evolution, floral development and chemical biosynthesis from the Aristolochia fimbriata genome. Nat. Plants 7, 1239–1253 (2021).

33. Y. Van de Peer, E. Mizrachi, K. Marchal, The evolutionary significance of polyploidy. Nat. Rev. Genet. 18, 411–424 (2017).

34. P. Ma et al., Genome assemblies of 11 bamboo species highlight diversification induced by dynamic subgenome dominance. Nat. Genet. 56, 710–720 (2024).

35. R. Zhang et al., Subgenome-aware analyses suggest a reticulate allopolyploidization origin in three Papaver genomes. Nat. Commun. 14, 2204 (2023).

36. P. Sun et al., Subgenome-aware analyses reveal the genomic consequences of ancient allopolyploid hybridizations throughout the cotton family. Proc. Natl. Acad. Sci. U. S. A. 121, e2313921121 (2024).

37. D. Wang et al., Ancient allopolyploidy and specific subgenomic evolution drove the radiation of poplars and willows. Nat. Commun. 16, 6881 (2025).

38. Y. H. Tseng, L. Y. Kuo, I. Borokini, S. Fawcett, The role of deep hybridization in fern speciation: Examples from the Thelypteridaceae. Am. J. Bot. 111, e16388 (2024).

39. M. Marcet-Houben, T. Gabaldón, Beyond the whole-genome duplication: phylogenetic evidence for an ancient interspecies hybridization in the baker’s yeast lineage. PLoS Biol. 13, e1002220 (2015).

40. J. F. Wendel, S. A. Jackson, B. C. Meyers, R. A. Wing, Evolution of plant genome architecture. Genome Biol. 17, 37 (2016).

41. P. P. Edger, M. R. McKain, K. A. Bird, R. VanBuren, Subgenome assignment in allopolyploids: challenges and future directions. Curr. Opin. Plant Biol. 42, 76–80 (2018).

42. A. M. Session, Allopolyploid subgenome identification and implications for evolutionary analysis. Trends Genet. 40, 621–631 (2024).

43. P. Sun et al., WGDI: A user-friendly toolkit for evolutionary analyses of whole-genome duplications and ancestral karyotypes. Mol. Plant 15, 1841–1851 (2022).

44. Y. Wang et al., MCScanX: A toolkit for detection and evolutionary analysis of gene synteny and collinearity. Nucleic Acids Res. 40, e49 (2012).

45. O. Garsmeur et al., Two evolutionarily distinct classes of paleopolyploidy. Mol. Biol. Evol. 31, 448–454 (2014).

46. Z. Guo et al., Deep learning can predict subgenome dominance in ancient but not in neo/synthetic polyploidized genomes. Plant J 10.1111/tpj.16979 (2024).

47. E. I. Alger, P. P. Edger, One subgenome to rule them all: underlying mechanisms of subgenome dominance. Curr. Opin. Plant Biol. 54, 108–113 (2020).

48. *Amborella* Genome Project, The *Amborella* genome and the evolution of flowering plants. Science 342, 1241089 (2013).

49. L. Zhang et al., The water lily genome and the early evolution of flowering plants. Nature 577, 79–84 (2020).

50. P. Sun et al., Early diversification and karyotype evolution of flowering plants. Research Square [Preprint] (2022). https://www.researchsquare.com/article/rs-1410884/v1 (accessed 22 June 2024).

51. L. Ma et al., Diploid and tetraploid genomes of Acorus and the evolution of monocots. Nat. Commun. 14, 3661 (2023).

52. Z. Wang et al., A high-quality *Buxus austro-yunnanensis* (Buxales) genome provides new insights into karyotype evolution in early eudicots. BMC Biol. 20, 216 (2022).

53. T. J. Shalev et al., The western redcedar genome reveals low genetic diversity in a self-compatible conifer. Genome Res. 32, 1952–1964 (2022).

54. Y. Liu et al., The *Cycas* genome and the early evolution of seed plants. Nat. Plants 8, 389–401 (2022).

55. F. A. Simão, R. M. Waterhouse, P. Ioannidis, E. V. Kriventseva, E. M. Zdobnov, BUSCO: Assessing genome assembly and annotation completeness with single-copy orthologs. Bioinformatics 31, 3210–3212 (2015).

56. H. Chen, A. Zwaenepoel, Y. Van de Peer, wgd v2: a suite of tools to uncover and date ancient polyploidy and whole-genome duplication. *Bioinformatics*, btae272 (2024).

57. M. Zhao, B. Zhang, D. Lisch, J. Ma, Patterns and consequences of subgenome differentiation provide insights into the nature of paleopolyploidy in plants. Plant Cell 29, 2974–2994 (2017).

58. F. Cheng et al., Gene retention, fractionation and subgenome differences in polyploid plants. Nat. Plants 4, 258–268 (2018).

59. Z. Liang, J. C. Schnable, Functional divergence between subgenomes and gene pairs after whole genome duplications. Mol. Plant 11, 388–397 (2018).

60. P. P. Edger et al., Subgenome Dominance in an Interspecific Hybrid, Synthetic Allopolyploid, and a 140-Year-Old Naturally Established Neo-Allopolyploid Monkeyflower. Plant Cell 29, 2150–2167 (2017).

61. L. Zeng et al., Resolution of deep angiosperm phylogeny using conserved nuclear genes and estimates of early divergence times. Nat. Commun. 5, 4956 (2014).

62. L. Cai et al., The perfect storm: Gene tree estimation error, incomplete lineage sorting, and ancient gene flow explain the most recalcitrant ancient angiosperm clade, malpighiales. Syst. Biol. 70, 491–507 (2021).

63. G. The Angiosperm Phylogeny, An update of the Angiosperm Phylogeny Group classification for the orders and families of flowering plants: APG III. Bot. J. Linn. Soc. 161, 105–121 (2009).

64. H. Hu, P. Sun, Y. Yang, J. Ma, J. Liu, Genome-scale angiosperm phylogenies based on nuclear, plastome, and mitochondrial datasets. J. Integr. Plant Biol. 65, 1479–1489 (2023).

65. C. Zhang, S. Mirarab, Weighting by gene tree uncertainty improves accuracy of quartet-based species trees. Mol. Biol. Evol. 39, msac215 (2022).

66. Y. Yang et al., Prickly waterlily and rigid hornwort genomes shed light on early angiosperm evolution. Nat. Plants 6, 215–222 (2020).

67. D. M. Emms, S. Kelly, OrthoFinder: Phylogenetic orthology inference for comparative genomics. Genome Biol. 20, 238 (2019).

68. S. Proost et al., i-ADHoRe 3.0—fast and sensitive detection of genomic homology in extremely large data sets. Nucleic Acids Res. 40, e11–e11 (2012).

69. K. Katoh, K. Misawa, K. i. Kuma, T. Miyata, MAFFT: a novel method for rapid multiple sequence alignment based on fast Fourier transform. Nucleic Acids Res. 30, 3059–3066 (2002).

70. C. Zhang, Y. Zhao, E. L. Braun, S. Mirarab, TAPER: Pinpointing errors in multiple sequence alignments despite varying rates of evolution. Methods Ecol. Evol. 12, 2145–2158 (2021).

71. B. Q. Minh et al., IQ-TREE 2: new models and efficient methods for phylogenetic inference in the genomic era. Mol. Biol. Evol. 37, 1530–1534 (2020).

72. J. C. Schnable, N. M. Springer, M. Freeling, Differentiation of the maize subgenomes by genome dominance and both ancient and ongoing gene loss. Proc. Natl. Acad. Sci. U. S. A. 108, 4069–4074 (2011).

73. T. Shi et al., Distinct expression and methylation patterns for genes with different fates following a single whole-genome duplication in flowering plants. Mol. Biol. Evol. 37, 2394–2413 (2020).

74. C. Zhang, R. Nielsen, S. Mirarab, ASTER: A package for large-scale phylogenomic reconstructions. *Mol. Biol. Evol.*, msaf172 (2025).

75. C. Zhang, C. Scornavacca, E. K. Molloy, S. Mirarab, ASTRAL-Pro: Quartet-based species-tree inference despite paralogy. Mol. Biol. Evol. 37, 3292–3307 (2020).

